# Genotype-free individual genome reconstruction of Multiparental Population Models by RNA sequencing data

**DOI:** 10.1101/2020.10.11.335323

**Authors:** Kwangbom Choi, Michael W. Lloyd, Hao He, Daniel M. Gatti, Vivek M. Philip, Narayanan Raghupathy, Mathew Vincent, Sai Lek, Isabela Gerdes Gyuricza, Steven C. Munger, Alan D. Attie, Mark Keller, Elissa J. Chesler, Karl W. Broman, Anuj Srivastava, Gary A. Churchill

**Author notes:** equal contribution (Kwangbom Choi, Michael W. Lloyd).

## Abstract

Multi-parent populations (MPPs), model organisms derived from two or more inbred founder strains, are widely used in biomedical and agricultural research. Gene expression profiling by direct RNA sequencing (RNA-Seq) is commonly applied to MPPs to investigate gene expression regulation and to identify candidate genes. In genetically diverse populations, including MPPs, quantification of gene expression is improved when the RNA-Seq reads are aligned to individualized transcriptomes that incorporate known polymorphic loci. However, the process of constructing and analyzing individual genomes can be computationally demanding and error prone. We propose a new approach, genome reconstruction by RNA-Seq (GBRS), that relies on simultaneous alignment of RNA-Seq reads to founder strain transcriptomes to reconstruct the diploid genome and quantify total and allele-specific gene expression in MPPs. We demonstrate that GBRS performs as well as methods that rely on high-density genotyping arrays. When used in conjunction with other genotyping methods, GBRS provides quality control for detecting sample mix-ups or contamination. GBRS software is freely available at https://github.com/churchill-lab/gbrs.

## INTRODUCTION

RNA sequencing (RNA-Seq) has revolutionized our understanding of gene expression in whole tissues and single cells (STARK et al. 2019). While primarily used to quantify transcript abundance, RNA-Seq data can also identify single nucleotide polymorphisms (SNPs) and small insertions and deletions (indels) in the transcribed genome (PISKOL et al. 2013), detect spontaneous mutations (MILLER et al. 2013), RNA editing events (GU et al. 2016), and quantify allele-specific expression (WITTKOPP et al. 2004). Here, we explore using RNA-Seq data for genotype reconstruction and individualized gene expression quantification in multi-parent populations (MPPs).

Typically, RNA-Seq analysis quantifies transcript abundance by counting reads aligned to a reference genome-based transcriptome index (FERRAGINA AND MANZINI 2000). Alignment algorithms can allow for mismatches due to sequencing errors and polymorphisms. However, reliance on a reference genome can introduce biases in gene expression quantification (DEGNER et al. 2009). These biases can be minimized by aligning RNA-Seq reads to a transcriptome that incorporates individual-specific genetic variants. We have shown that alignment to individualized transcriptomes significantly improves expression quantitative trait locus (eQTL) mapping (MUNGER et al. 2014). This approach requires prior knowledge of individual genomes, obtained through whole genome sequencing or marker genotyping and variant imputation. Constructing alignment indices for each sample can be error-prone if individual genome reconstructions are not accurate.

MPPs are genetic reference populations derived from two or more inbred founder strains (DE KONING AND MCINTYRE 2017). Each MPP individual’s genome is a mosaic of founder strain genome segments. For many MPPs, the founder strains’ whole genome sequences have been assembled and annotated. MPP individuals can be genetically characterized using genotyping arrays (MORGAN et al. 2015) or short-read DNA sequencing (PARKER et al. 2016) to detect known founder strain variants. The genome mosaic can be reconstructed using a Hidden Markov Model (HMM) (GATTI et al. 2014; BROMAN et al. 2019). The full diploid genome sequence of an MPP individual can then be inferred by imputing variants onto the founder haplotype blocks of the mosaic MPP genome (MUNGER et al. 2014).

In any tissue, thousands of expressed genes are distributed across the genome. Variants detected by RNA-Seq could potentially replace genotyping arrays. However, it is unclear if RNA-Seq data capture enough information for accurate genome reconstruction. Direct approaches based on variant calling from RNA-Seq reads require deep sequencing coverage (LOPEZ-MAESTRE et al. 2016) and may not be reliable, especially for genes with low expression. Additionally, the distribution of expressed genes may not be dense or uniform enough to accurately reconstruct haplotypes in some regions. Here we propose a novel solution, GBRS (<u>G</u>enome reconstruction <u>By</u> <u>R</u>NA-<u>S</u>eq), to quantifying gene expression from MPPs. GBRS avoids the step of creating individual genomes by employing a multi-way alignment index that represents the combined transcriptomes of the MPP founder strains. We demonstrate that the individual genome reconstruction and gene expression quantification from GBRS perform as well as methods that rely on high-density genotyping arrays. GBRS is implemented in an open-source Python package available at https://churchill-lab.github.io/gbrs/, and wrapped into Nextflow (DI TOMMASO *et al*. 2017) pipelines available at https://github.com/TheJacksonLaboratory/cs-nf-pipelines.

### Overview of GBRS algorithm

The objectives of GBRS are to 1) reconstruct the founder haplotype mosaic of individuals from a multi-parent population (MPP) directly from RNA-Seq data and 2) quantify total and allele-specific gene expression based on individual MPP genomes. GBRS uses a multi-way alignment index built with the combined predicted transcript sequences of the MPP founder strains. At each gene locus the multi-way index represents all predicted transcript isoforms for each founder strain. We hypothesized that the specificity of RNA-Seq read alignments to the multi-way index would provide information about local (gene-level) genotypes. Since individual genes may not have sufficient sequence variation to uniquely identify a genotype, we employ a Hidden Markov Model (HMM) to combine information across neighboring genes to reconstruct the diploid mosaic of founder haplotypes that make up the genome of each MPP individual. Finally, we quantify gene expression based on read alignment to the diploid genome reconstruction.

When RNA-Seq reads are aligned to a multi-way index, many reads will align identically to multiple founder transcripts depending on the number and strain distribution pattern of variants spanned by the read. To resolve these multi-mapped reads, we compute the expected read counts for each founder transcript using the allele-specific weighted allocation algorithm EMASE (RAGHUPATHY et al. 2018). When the RNA-Seq reads come from one of the founder strains, the read counts will exhibit characteristic proportions that we refer to as the *founder profile* of a gene. The founder profile is a vector of proportions of read counts allocated to each founder in the multi-way index. It indicates how reads from a known founder strain align to their own predicted transcripts as well as to transcripts of other founder strains. In general, the more variants that distinguish the founder strain transcripts, the more specific the founder profiles will be. For a gene with multiple variants that have distinct strain distribution patterns, the matrix formed by combining all founder profiles will approach one on the diagonal elements. For a gene with no variants, the proportions in each founder profile will be equal, reflecting our inability to distinguish the founder strain origin of RNA-Seq reads from this gene.

Founder profiles depend only on predicted transcript sequences and RNA-Seq reads from founder strain samples. Thus, we hypothesized that the same profile will be reproduced in read counts from an MPP individual that is homozygous for the founder haplotype at a gene. We note that this should be true even if the predicted founder strain transcript is not accurate. We further assume that read counts at a heterozygous locus will present as a mixture of the two corresponding founder profiles.

To apply GBRS to an individual MPP sample, we first align the RNA-Seq reads from the individual to the multi-way alignment index and compute the *sample profiles*—the proportion of reads from an individual sample that align to each of the founder transcripts at a given gene (**Figure 1a**). The observed sample profiles are analyzed using a hidden Markov model (HMM) to estimate the genotype state of the MPP sample across expressed genes (**Figure 1b**). The HMM has two components: an emission model and a transition model. The emission model describes the probability distribution of a sample profile (*y*_*t*_) for a given genotype state (*s*_*t*_) at a gene locus *t*. The founder profile (*x*_*t*,#_) defines the center of the emission probability distribution for an assumed genotype state *g*. The emission model is based on the multivariate normal distribution:

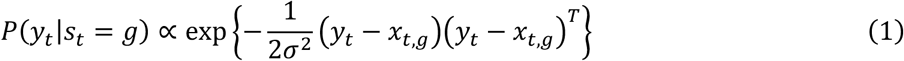

**Figure 1:**
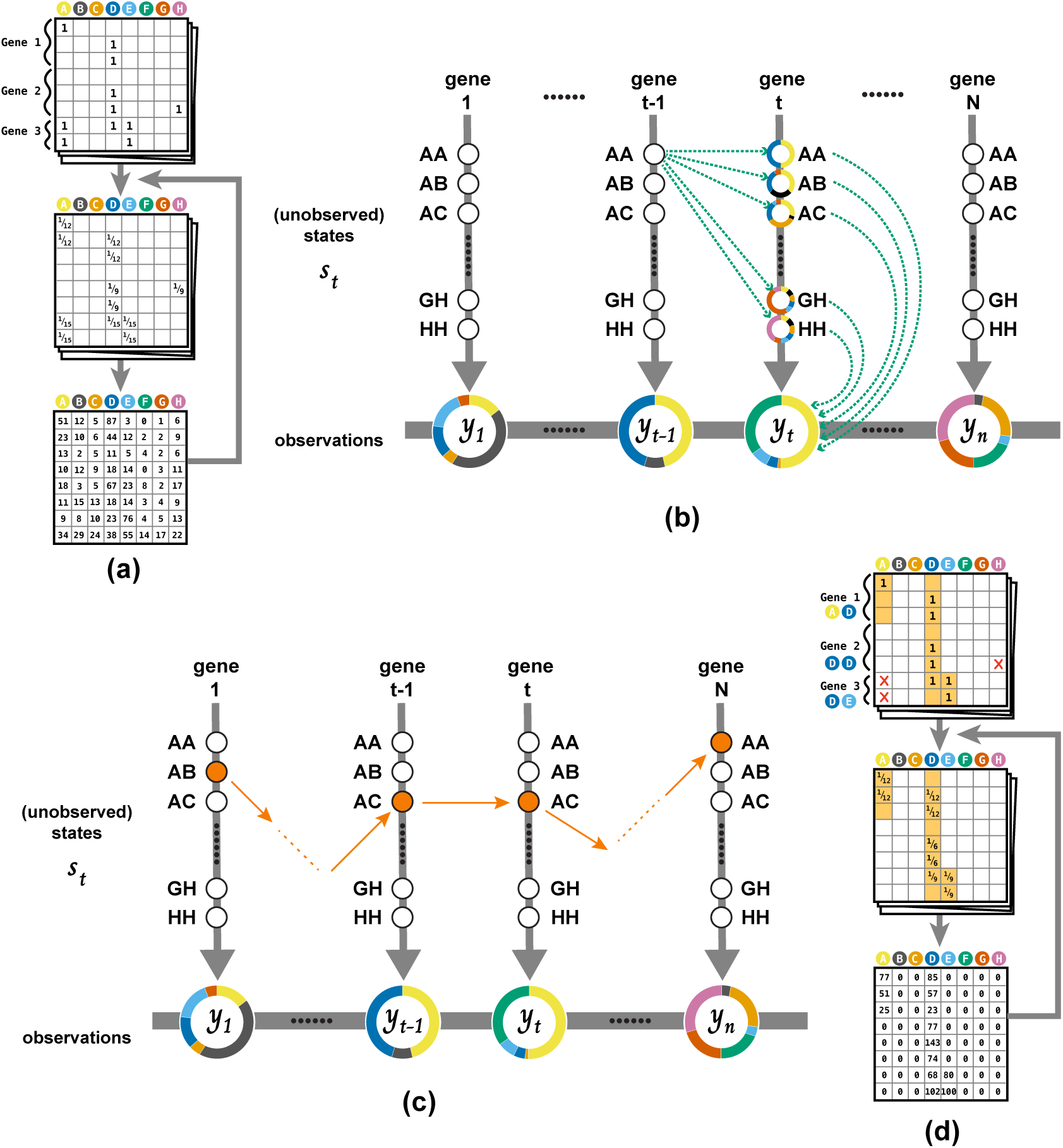
Overview of the GBRS algorithm. (**a**) The multiway alignment index is composed of all annotated gene transcripts across the eight founder strains. The incidence matrix for a single RNA-Seq indicates where identically best alignments occur. For multi-mapped reads, the incidence matrix is reallocated with weights proportional to the probability of the alignment location. The weights are summed across reads to obtain estimated expected read counts, and the process is iterated to convergence. The estimated read counts for across all transcripts of each gene are summed and converted to proportions to obtain sample profiles. (**b**) The HMM architecture has 36 ‘hidden’ genotype states (AA, AB, …) at each gene, one of which has given rise to the corresponding sample profile (circles, *yt*). The transition model defines the probability of changes in genotype states between genes t-1 and t (straight arrows), and the emission model measures the similarity between the sample profile and the pre-computed founder profiles (curved arrows). A single pass forward-backward algorithm uses the sample profiles to compute the marginal probability of the genotype state at each gene. (**c**) The Viterbi algorithm is applied to the genotype probabilities to compute the most probable sequence of genotype states across all gene loci – the genome reconstruction. (**d**) We return to the read alignment incidence matrix and mask read alignments that do not correspond to the genome reconstruction. We repeat the weighted allocation to obtain diploid allele-specific expression estimates.

where σ^2^ is a tuning parameter that we set to 0.12, a value that approximates the theoretically expected number of recombination events (GATTI *et al*. 2014). The transition model describes how the genotype states can change across the intervals between genes. The transition probability, *P*(*s*_*t*_|*s_t_*_-’_), is a function of the breeding history and the distance between neighboring gene loci *t* − 1 and *t*. We analyze each MPP individual to compute the posterior probability of genotypes at each gene locus, *P*(*s_t_*|*Y*), with a one-time execution of the forward-backward algorithm (RABINER 1989).

To obtain a maximum-probability reconstruction of the diploid MPP genome, we apply a Viterbi algorithm (VITERBI 1967) to the genotype probabilities to find the most likely diploid genotype at each gene (**Figure 1c**). To estimate allele-specific expression, we repeat the weighted allocation of the MPP RNA-Seq reads with the restriction that reads align only to the founder transcripts in the inferred diploid genotype (RAGHUPATHY *et al*. 2018) (**Figure 1d**). The reallocation step ensures that the allele-specific counts reflect the most probable diploid MPP genome, and it does not require re-alignment of the RNA-Seq reads, so it is fast.

## METHODS

### Building the multi-way alignment index

We created custom genomes of the founder strains by introducing strain-specific SNPs and short indels from the Mouse Genome Project (KEANE *et al*. 2011) release v8 (https://ftp.ebi.ac.uk/pub/databases/mousegenomes/REL-2112-v8-SNPs_Indels/) to the reference genome GRCm39 Ensembl v105 primary assembly (http://ftp.ensembl.org/pub/release-105/fasta/mus_musculus/dna) using g2gtools (http://churchill-lab.github.io/g2gtools). We adjusted the coordinates of the reference gene annotation (GTF) Ensembl Release 105 (HOWE *et al*. 2021) (http://ftp.ensembl.org/pub/release-105/gtf/mus_musculus/) and extracted strain-specific transcripts with g2gtools. We filtered the GTF to include only the following biotypes: protein_coding, lncRNA, IG_C_gene, IG_D_gene, IG_J_gene, IG_LV_gene, IG_V_gene, TR_C_gene, TR_D_gene, TR_J_gene, and TR_V_gene. We appended the founder strain codes (e.g., A, B, C, *· · ·*, H for A/J, C57BL/6J, 129S1/SvImJ, NOD/ShiLtJ, NZO/HILtJ, CAST/EiJ, PWK/PhJ, and WSB/EiJ, respectively) to each transcript ID; collated the founder transcript sequences into a single fasta file, and ran the bowtie-build from Bowtie v1.3.1 (LANGMEAD *et al*. 2009).

### Training the HMM

We obtained RNA-Seq data from the DO founder strains across six tissues (62 adipose, 61 gastrocnemius, 63 heart, 64 hippocampus, 189 liver, and 95 pancreatic islet). In total, we used 534 inbred mice (266 male and 268 female) (**Table 1**). We aligned RNA-seq data to the multi-way index using Bowtie v1.3.1 (LANGMEAD *et al*. 2009) with ‘all’, ‘best’, and ‘strata’ options. Many reads mapped to multiple predicted founder transcripts of a gene as well as to multiple genes. We transformed the raw read alignment counts to estimated expected counts using weighted allocation by expectation maximization for allele-specific expression (EMASE) (RAGHUPATHY *et al*. 2018). We detected 15,802 genes with mean expression above 1 transcript per million (TPM). Then we converted the expected counts to proportions to obtain the estimated founder profiles. This one-time training process provides estimated founder profiles (*x*_*t*,#_) for each founder strain *g* at each gene position *t*.

**Table 1.** Founder inbred strain samples used to estimate the founder profiles.

| Strain | Sex | Adipose | Gastrocnemius | Heart | Hippocampus | Liver | Pancreatic Islet |
| --- | --- | --- | --- | --- | --- | --- | --- |
| A/J | Female | 4 | 4 | 4 | 4 | 12 | 7 |
|  | Male | 3 | 4 | 4 | 4 | 11 | 7 |
| C57BL/6J | Female | 4 | 4 | 4 | 4 | 12 | 9 |
|  | Male | 4 | 4 | 4 | 4 | 11 | 9 |
| 129S1/SvImJ | Female | 4 | 4 | 4 | 4 | 12 | 5 |
|  | Male | 4 | 4 | 4 | 4 | 12 | 5 |
| NOD/ShiLtJ | Female | 4 | 4 | 4 | 4 | 12 | 5 |
|  | Male | 4 | 3 | 4 | 4 | 12 | 5 |
| NZO/HILtJ | Female | 4 | 4 | 4 | 4 | 12 | 3 |
|  | Male | 4 | 4 | 4 | 4 | 12 | 1 |
| CAST/EiJ | Female | 3 | 3 | 3 | 4 | 11 | 6 |
|  | Male | 4 | 4 | 4 | 4 | 12 | 8 |
| PWK/PhJ | Female | 4 | 4 | 4 | 4 | 12 | 4 |
|  | Male | 4 | 3 | 4 | 4 | 12 | 5 |
| WSB/EiJ | Female | 4 | 4 | 4 | 4 | 12 | 9 |
|  | Male | 4 | 4 | 4 | 4 | 12 | 7 |
|  | Sum | 62 | 61 | 63 | 64 | 189 | 95 |

We obtained the theoretical transition probability according to (BROMAN 2005; BROMAN 2012) using genetic distances between the start sites of neighboring genes. We obtained the transition probabilities for each breeding generation of DO mice using DOQTL (GATTI *et al*. 2014) v1.17.5.

### Applying GBRS to MPP Data

We applied GBRS to RNA-sequencing data from 483 DO mice (KELLER et al., 2018; TYLER et al., 2024), equally divided between male and female, from breeding generations 17 to 23. We profiled five tissues (469 adipose, 483 gastrocnemius, 482 heart, 482 liver, and 378 pancreatic islet) for a total of 2,294 RNA-seq samples (**Table 2**).

**Table 2.** Diversity outbred (DO) sampled used in this study for demonstration of GBRS and downstream eQTL mapping.

| Sex | Adipose | Gastrocnemius | Heart | Liver | Pancreatic Islet |
| --- | --- | --- | --- | --- | --- |
| Female | 228 | 240 | 239 | 240 | 188 |
| Male | 241 | 243 | 243 | 242 | 190 |
| Total | 469 | 483 | 482 | 482 | 378 |

Raw reads were aligned with Bowtie v1.3.1 (LANGMEAD *et al*. 2009) to the multi-way index. Paired-end reads (adipose, gastrocnemius, heart, liver) were mapped individually, converted to EMASE format (GBRS *compress*), and re-paired (EMASE *get-common-alignment*). For single-end data (pancreatic islet), read were mapped individually and converted to EMASE format (GBRS *compress*). Genotype reconstruction used the HMM described above (GBRS *reconstruct*). Diploid expression profiles (GBRS *quantify-genotypes*), and genotype probability matrices are inferred (GBRS *interpolate*). Expression was then quantified using the EMASE algorithm (GBRS *quantify*). All analysis was conducted with GRBS v1.0.0 (https://github.com/churchill-lab/gbrs).

### Comparison of GBRS Genotypes to Genotyping Arrays

To compare genotype reconstruction from GBRS against array-based genotyping, the 483 DO mice used in RNA sequencing were also genotyped on the GigaMUGA mouse universal genotyping array which contains *∼*143,000 markers distributed throughout the mouse genome (GeneSeek, Lincoln, NE). The GigaMUGA genotypes were based on tail-tip samples collected for DNA analysis (KELLER et al, 2018). Low-quality GigaMUGA results with greater than 10% missing genotypes were removed, leaving 455 adipose, 469 muscle, 468 heart, 469 liver, and 364 pancreatic islet samples that had both GBRS and GigaMUGA genotyping data. The genotype probabilities for the GigaMUGA were obtained using the DO-specific HMM (BROMAN 2009) implemented in the R/qtl2 package (BROMAN et al., 2019). To facilitate comparisons, we interpolated the genotype probabilities from both methods onto a common, evenly spaced grid of 74,165 pseudo-markers.

To evaluate the agreement of genome reconstructions between GBRS and the genotyping array, we collapsed the 36-state genotype probabilities to 8-state haplotype dosages (GATTI *et al*. 2014) and computed the Pearson correlation of the vectorized haplotype dosages. We assumed that a sample was mismatched between the GBRS and GigaMUGA if the correlation fell below 0.8 For each mismatched sample, we then searched for a sample match by comparing the GBRS to all available GigaMUGA samples and vice-versa.

We computed the number of predicted recombination events, as well their location and flanking founder strain haplotypes, based on the most-probable diploid genome reconstruction (Viterbi path) for both the GBRS and GigaMUGA array data.

### eQTL analysis

We mapped eQTL using sample corrected GBRS genotypes and gene-level expected read counts estimated by GBRS. Genes with median of count value *>* 1 TPM were included in the eQTL analysis. Raw counts in each sample were normalized with the variance-stabilizing transformation (VST) in the DESeq2 R package (LOVE *et al*. 2014). A linear mixed model with sex, diet and generation as additive covariates and a random polygenic term to account for genetic relatedness was fit at each locus on the pseudo-marker grid using *qtl2* R package (BROMAN et al., 2019).

Significance thresholds were established by performing 1,000 permutations and fitting an extreme value distribution to the genome-wide maximum LOD scores (DUDBRIDGE AND KOELEMAN 2004). Permutation derived P-values were then converted to a false discovery rate (FDR, q-value) across expressed genes with the *qvalue* R package (STOREY *et al*. 2020), using the bootstrap method to estimate *π*_0_ and the default *λ* tuning parameters (STOREY *et al*. 2003). We detected QTL at the adjusted genome-wide significance level of FDR < 0.05.

### Simulated MPP Data

We constructed a synthetic expression profile based on expected RNA-Seq read counts estimated from a randomly selected DO tissue sample using GBRS. We used this synthetic RNA-Seq data to compare GBRS RNA expression quantification to another widely used method -RSEM (LI AND DEWEY 2011) that relies on read alignment to the mouse reference genome (GRCm39). To generate the synthetic expression data, we assigned diploid DO genotypes from GBRS to the filtered GRCm39 Ensembl transcript set described above. We set a “true” TPM count for each transcript based on a random sample of the exponential distribution. Heterozygous transcripts were assigned equal TPM for each allele. We used the R package *Rsubread* (LIAO *et al*. 2019) to simulate 30 million 2×100bp paired-end reads (min fragment length: 125, max fragment length: 500, mean fragment length: 250, fragment length standard deviation: 40) using the randomly simulated TPM values and allele-specific transcript sequences. The simulated RNA-Seq data was analyzed using GBRS as described above. We also aligned the RNA-Seq reads to the reference genome using Bowtie with the same settings as for GBRS except that the paired-end data were mapped in PE mode. Reads mapped to GRCm39 were quantified by RSEM v1.3.1. The GBRS expression quantification was then compared to synthetic true values and to RSEM quantified expression.

## RESULTS

### Diversity Outbred Mapping and Quantification Summary

We obtained RNA-Seq data across six tissues from a cohort of 483 DO mice and aligned the sequenced reads to a multi-way alignment index. Alignment percentages ranged from 58 to 81% of raw reads (adipose: mean 80.7, SD 3.99; gastrocnemius: mean 81.2, SD 2.72; heart: mean 58.2, SD 6.46; liver: mean 80.1, SD 3.91; pancreatic islet: mean 74.9, SD 6.68). We detected between 11,166 and 13,684 genes per tissue after removing genes with average expression below 1 TPM (adipose: 13,684; gastrocnemius: 11,242; heart: 11,849; liver: 11,166; pancreatic islet: 12,608). Intersecting the detected expressed genes in our DO samples with the 15,802 genes that had estimated founder profiles in the GBRS HMM left 13,009 adipose, 11,052 gastrocnemius, 11,620 heart, 11,038 liver, and 12,033 pancreatic islet genes for further analysis.

### GBRS produces accurate haplotype reconstructions

An array of genotype probabilities is a key data object used for QTL mapping, variant imputation, and other downstream genetic analyses of MPP populations (BROMAN *et al*. 2019). The array has dimensions of samples*×*markers*×*genotypes, and sums to one across genotypes for each sample and marker. The genotype probabilities capture uncertainty in estimation of the pair of founder haplotypes present in each MPP individual at each marker locus. For DO mice, with eight founder strains, there 36 possible diploid genotypes which can be reduced to eight haplotype dosages (GATTI *et al*. 2014). We compared genotype probabilities estimated by GBRS to those estimated from the GigaMUGA arrays by computing the Pearson correlation of estimated haplotype dosages. We repeated this comparison using GBRS estimates from each tissue and found that more than 97% of all DO samples had Pearson correlation *r >* 0.8 (**Figure 2**). The median correlation coefficient for each tissue was 0.93 for adipose, 0.92 for gastrocnemius, 0.93 for heart, 0.92 for liver, and 0.93 for pancreatic islet. Next, we investigated the small proportion of samples that showed low correspondence between GBRS and GigaMUGA genotypes.

**Figure 2:**
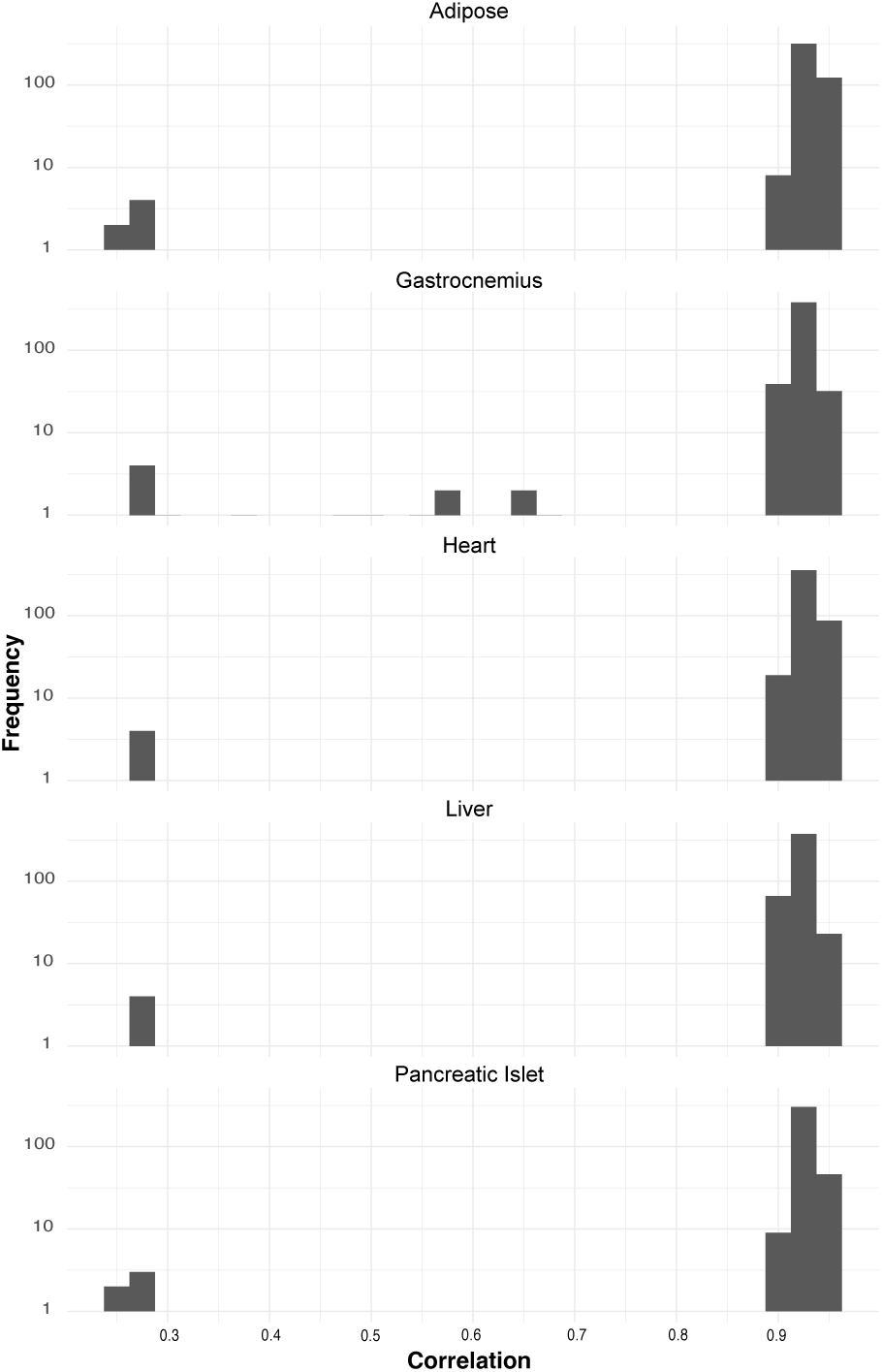
Correlation of GBRS and array-based genotype probabilities. Individual genomes reconstructed by GBRS are concordant with GigaMUGA reconstructions (*r >* 0.8) for the majority of MPP samples across tissues. The median of Pearson correlation of genotype probabilities is 0.93 for adipose; 0.92 for gastrocnemius (skeletal); 0.93 for heart; 0.92 for liver; and 0.93 for pancreatic islet. Nine samples show low Pearson correlation, indicating that the DNA and RNA samples do not correspond due to possible mix ups in sample handling.

### GBRS can identify and correct sample mix-ups

In studies that generate large numbers of tissue samples for multiple assays, one should always keep in mind the possibility of sample mix-ups (BROMAN *et al*. 2015). In our comparison of GBRS and GigaMUGA haplotype reconstructions, we identified six mice with discordant results (Pearson correlation *r* < 0.3). We compared the GBRS genotypes of these samples to all available GigaMUGA genotypes, and vice-versa. These comparisons identified four individuals in two pairwise swaps that occurred across all tissues. Within adipose there was one additional pairwise swap that occurred only within the adipose tissue. We conclude that the shared sample swaps were due to mislabeling of the DNA samples that were run on the GigaMUGA array, and the isolated swap was due to mislabeling of two adipose RNA samples. Thus, we were able to identify and correct these sample labeling errors. We found an additional nine samples within gastrocnemius with correlations between 0.3 and 0.8. We were not able to identify matching GigaMUGA genotypes for these samples and conclude that the low correlations were likely due to problems with the RNA sample quality. In subsequent analyses, we corrected the identifiable sample swaps and excluded the nine questionable samples from further analysis.

### GBRS accurately detects recombination events

To reconstruct founder haplotype mosaic of the MPP individual genomes, we computed the most probable diploid genotypes at each pseudo-marker locus by applying the Viterbi algorithm to the genotype probabilities estimated using GBRS and GigaMUGA methods. We localized recombination events to intervals between markers where the flanking marker genotypes were different (**Figure 3**). In an outbreeding MPP such as the DO, recombination breakpoints accumulate at a predictable linear rate with each breeding generation (GATTI *et al*. 2014). We assessed whether GBRS could detect recombination breakpoints with the same sensitivity as the MUGA genotyping arrays by comparing the numbers of detected events in each generation (G17-G23) of DO mice. GBRS detected on average, between 0.5% fewer and 5.82% more breakpoints depending on tissue and generation (**Table 3**; **Figure 4**). We conclude that GBRS reconstructions based on ∼11 to 13 thousand expressed genes are comparable to the high density (∼143k markers) GigaMUGA platform in DO mice from these outbreeding generations.

**Figure 3:**
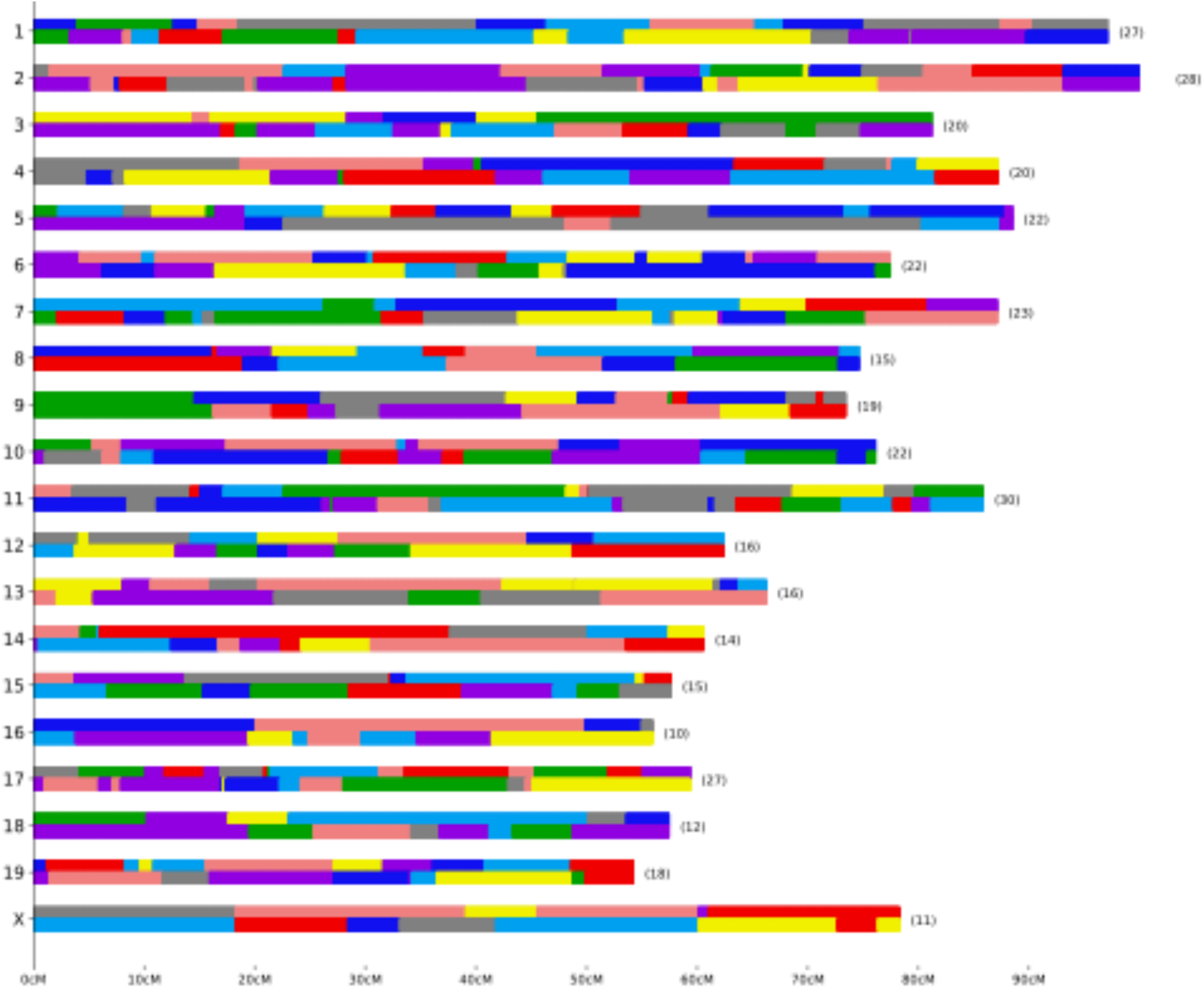
A Viterbi reconstruction of the haplotype mosaic of an individual MPP genome. GBRS reconstructed the diploid genome for a female liver sample set and identified 387 recombination breakpoints. The founder origins of haplotype blocks in indicated by color coding.

**Table 3.** Percent change in number of crossover events between GBRS and GigaMUGA array based estimates. Positive values indicate GBRS has called more events than GigaMUGA.

| Generation | Adipose | Gastrocnemius | Heart | Liver | Pancreatic Islet |
| --- | --- | --- | --- | --- | --- |
| 17 | 4.97 | 1.05 | 2.19 | 1.93 | 3.50 |
| 18 | 4.85 | 0.73 | 0.81 | 1.56 | 2.81 |
| 19 | 5.33 | 0.71 | 0.75 | 0.96 | 1.91 |
| 21 | 3.64 | -0.92 | 0.15 | 0.45 | 0.59 |
| 23 | 5.82 | -0.49 | 0.50 | 0.45 | NA |

**Figure 4:**
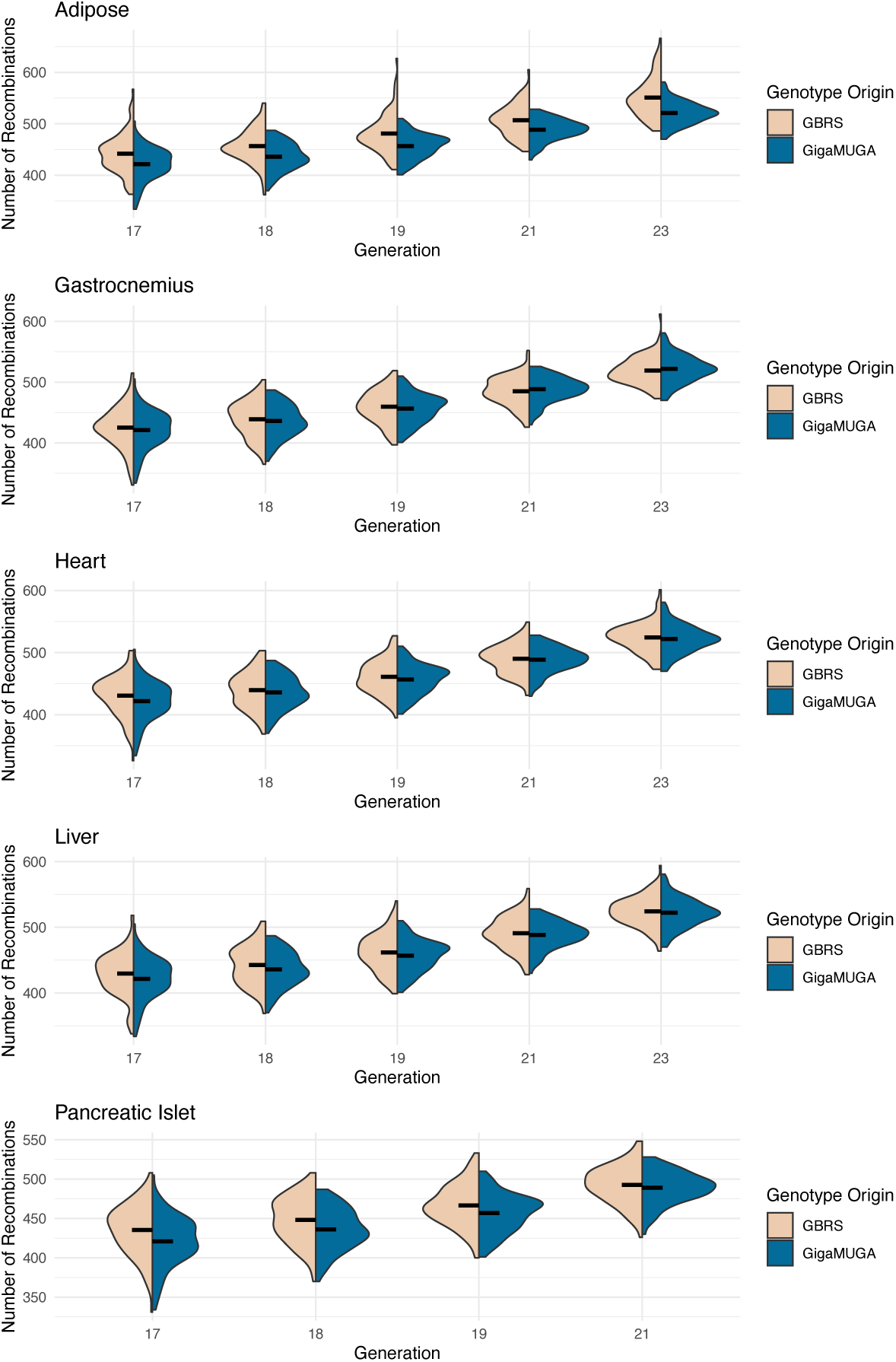
Estimated genome-wide recombination events. Comparision of the distribution of predicted recombination breakpoint counts predicted by GBRS (tan) and from GigaMUGA (blue) array data for all Diversity Outbred (DO) samples across adipose, gastrocnemius, heart, liver and pancreatic islet as a function of DO generation.

### GBRS genotypes support robust expression QTL mapping

Thousands of genes have expression levels that are influenced by genetic variation near the location of their coding sequences (local eQTL) or other genomic loci (distal eQTL) (CHICK *et al*. 2016; AGUET *et al*. 2017). These genetic associations can be by mapping gene expression levels, and local eQTL are more prevalent and typically show stronger association than distal eQTL. Incorrect genotypes are likely to reduce the strength of genetic association for both types of eQTL. We compared the strength of association at local and distal eQTL between GBRS genotypes and GigaMUGA genotypes by comparing the LOD scores from for all eQTL peaks detected at genome-wide significance (FDR < 0.05) for either genotyping method (**Figure 5**). For local eQTL, we found 6003, 6522, 6906, 5945, and 6611 genes with higher LOD scores and 5322, 5893, 5985, 5378, and 6300 genes with lower LOD scores in adipose, gastrocnemius, heart, liver, and pancreatic islet, respectively, using GBRS in comparison to GigaMUGA. For distal eQTL, we found 3047, 2474, 2122, 1995 and 4642 genes with higher LOD scores and with 4959, 3509, 2584, 2717, and 5872 genes have lower LOD scores using GBRS. Thus, while GBRS delivered slightly higher LOD scores for local eQTL and slightly lower LOD scores for distal eQTL, the two sets of eQTL mapping results are strikingly concordant.

**Figure 5:**
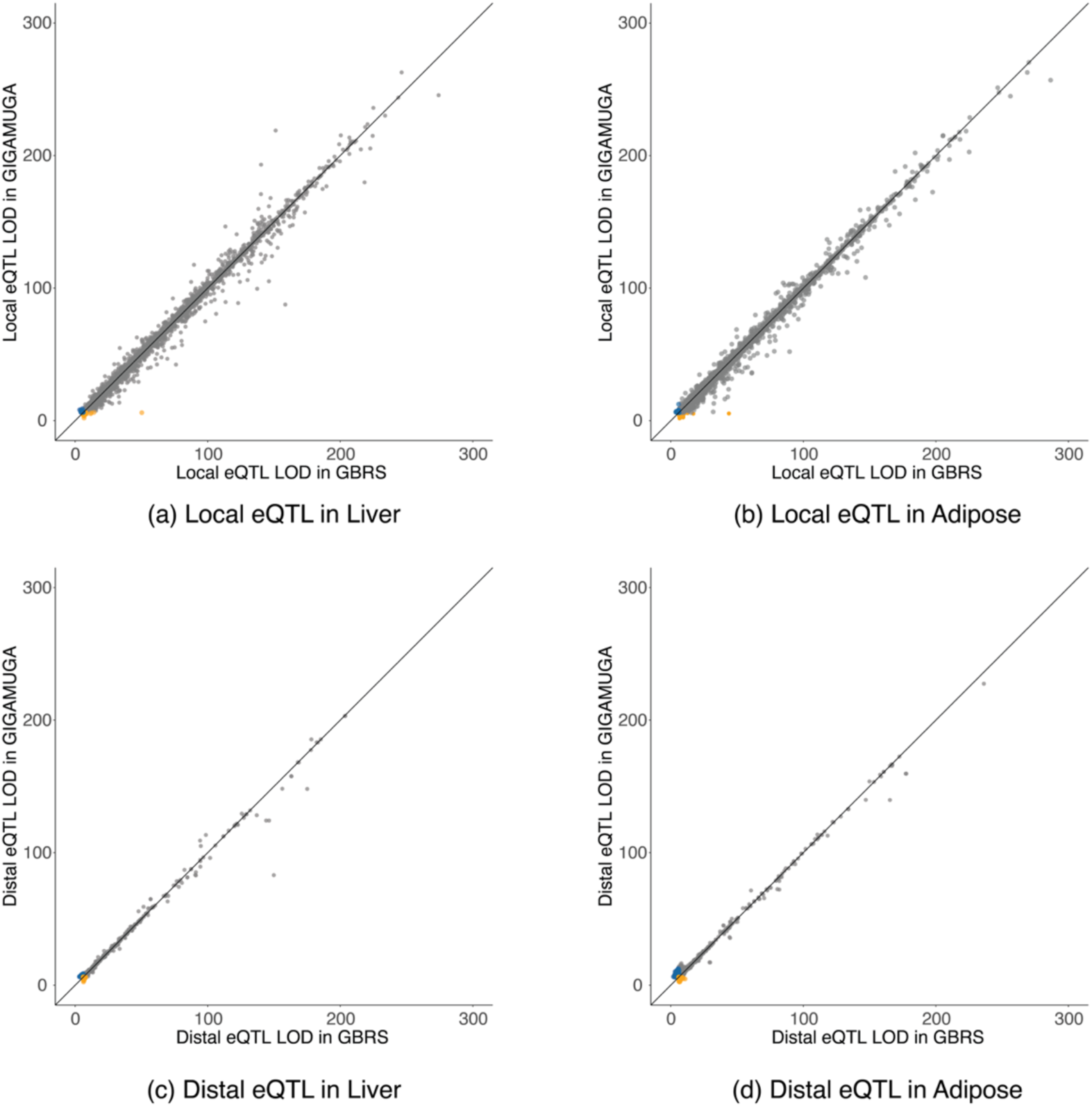
eQTL mapping analysis. LOD scores for eQTL obtained with GBRS and GigaMUGA array genotype probabilities. The LOD scores greater than 6 of local eQTL in (**a**) liver, (**b**) adipose, and distal eQTL in (**c**) liver, and (**d**) adipose tissues are shown as scatterplots. Orange points indicate significant LOD scores for GBRS over GigaMUGA, and blue points indicate significant scores for GigaMUGA over GBRS. An identity line is drawn on each scatterplot for reference.

### Simulated MPP Expression Profile Comparison

We simulated 30 million paired-end reads as described in METHODS, of which 86.5% aligned to the multi-way transcriptome. We examined the concordance of read counts estimated by GBRS with simulated ‘truth’ at the gene level (total), the transcript level, and the allele-specific transcript level. For total gene expression, GBRS was highly concordant with simulated TPM values (*R^2^* = 0.999; **Figure 6a**). At the total transcript level GBRS was slightly less efficient (*R^2^* = 0.97; **Figure 6b**). When examining simulated expression of individual transcript alleles, although GBRS failed to quantify some transcripts correctly the concordance was still quite high (*R^2^* = 0.91; **Figure 6c**). We further note that GBRS assigned 112,989 of 118,037 genotypes correctly (95%).

**Figure 6:**
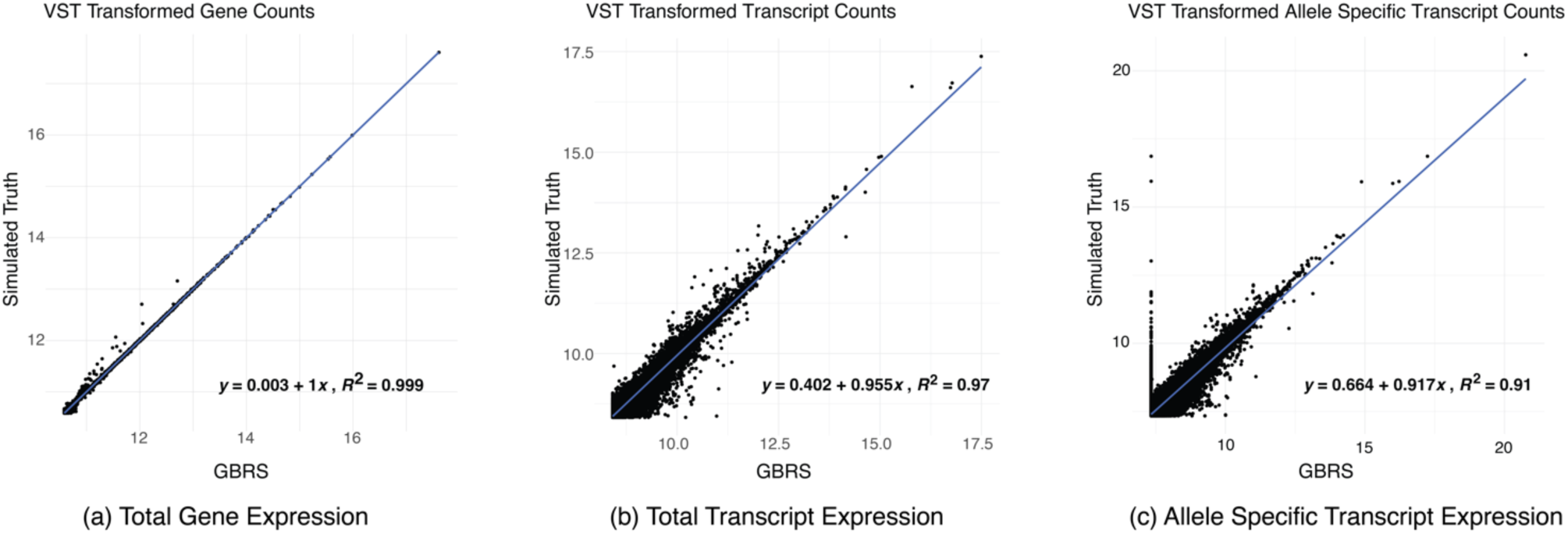
Analysis of simulated RNA-Seq data. GBRS quantified raw counts versus the simulated ‘truth’ counts for (**a**) total gene expression, (**b**) total transcript expression and (**c**) allele specific transcript expression.

Next, we compared GBRS to RSEM gene expression quantification. We mapped 70.6% of reads to the reference genome using Bowtie (METHODS). We applied RSEM for expression quantification to replicate a standard approach for this data type. In comparing GBRS with RSEM based expression quantification, both were comparable for both gene (*R^2^* = 0.99; **Figure 7a**) transcript level (*R^2^* = 0.95; **Figure 7b**) expression. We are unable to compare allele specific expression, as RSEM does not provide such estimates.

**Figure 7:**
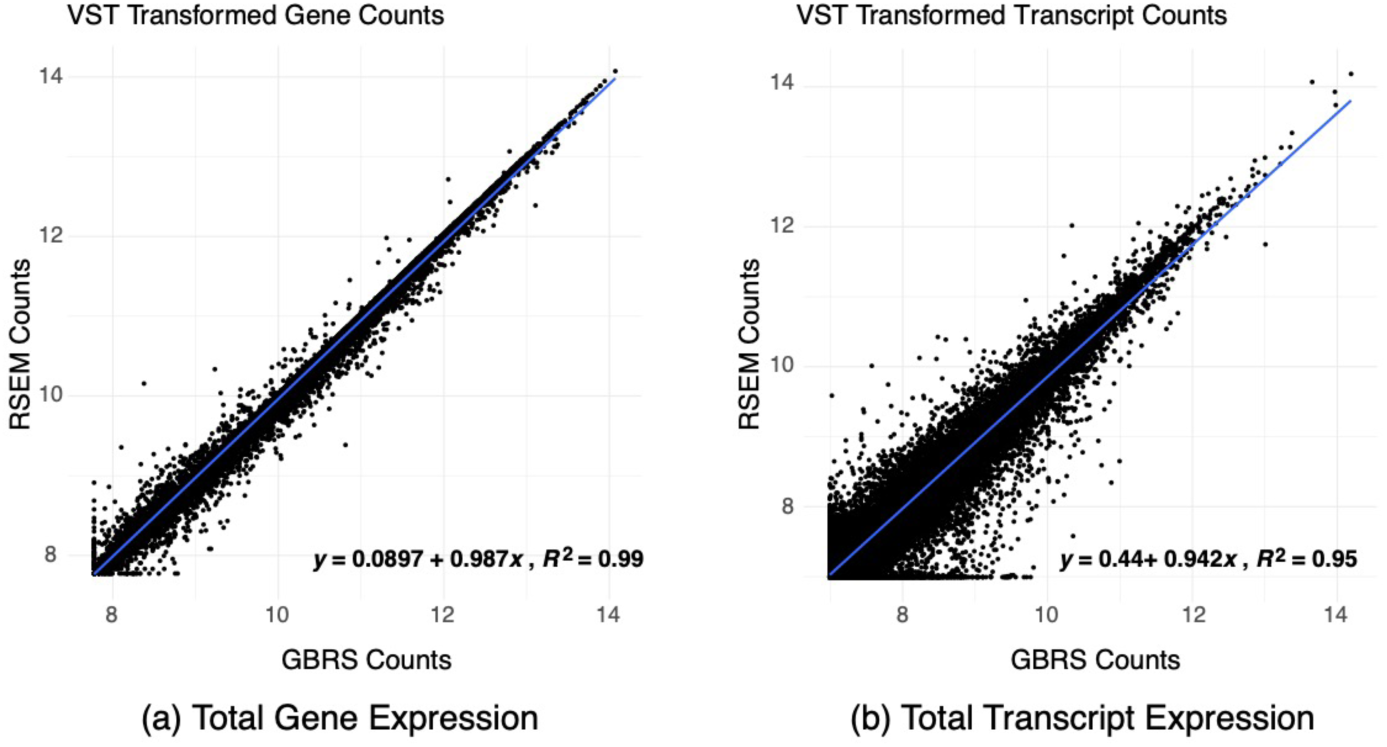
Comparison of GBRS with RSEM. GBRS quantified raw counts versus RSEM quantified counts for (**a**) total gene expression, and (**b**) total transcript expression.

## DISCUSSION

GBRS reconstructs the individual genomes of multi-parent population (MPP) individuals from RNA-Seq data and simultaneously quantifies total and allele-specific gene expression. It uses the specificity of multi-way RNA-Seq read alignment to founder transcripts and combines information from flanking genes to estimate the genotype at each expressed gene locus using probabilistic genotypes to capture uncertainty. GBRS also provides a maximum probability estimate of the genome-wide haplotype mosaic that constitutes and MPP individual genome. Unlike other genotyping strategies that use sequencing data, GBRS does not rely on variant calling. GBRS can deliver the advantages of aligning RNA-Seq reads to individual genomes (MUNGER *et al*. 2014) without the need to construct many individual alignment indices.

GBRS treats the gene as a unit and does not account for recombination within a gene. Moreover, the distribution of genes is uneven across the genome, which may limit the resolution of mapping breakpoints. It may be possible to improve resolution by estimating genotype probabilities at individual exons, but precision will still be limited by the number and location of expressed genes. As the DO mice, and other outbred MPPs evolve, the density of recombination breakpoints will eventually increase beyond the resolution of GBRS genotype reconstructions.

In the multi-way alignment index, a gene is represented by the collection of all annotated transcript isoforms for each founder strain. Isoforms will often share high levels of sequence identity within and between founder strains, which results in a high proportion of multi-mapping reads. Weighted allocation algorithms (LI et al., 2011; RAHGUPATHY et al., 2018) resolve ambiguity in read mapping but they are more reliable when there is less redundancy. Aggregated read counts are more accurate. We obtained the highest accuracy for estimates of total gene expression (summed across transcripts and alleles) but estimates of transcript-level and allele-specific transcript-level expression also agree with simulated truth and in comparisons to RSEM (all R^2^ > 0.9).

GBRS seems well suited for eQTL mapping because the genotypes are estimated at the locations of the expressed genes. We were concerned that local eQTL LOD scores might be inflated. However, the differences in GBRS versus GigaMUGA LOD scores (**Figure 5**) suggest that any bias is small.

To ‘train’ the GBRS algorithm, we used founder strain RNA-Seq samples to estimate founder profiles in the HMM emission model. We assumed that the sample profiles at a homozygous gene locus would match the corresponding founder profile. We further assumed that a heterozygous sample profile would look like an equally weighted mixture of two founder profiles, which may not be accurate if there is allelic imbalance of gene expression. Training with F1 hybrid samples was not practical. If founder RNA-Seq data had not been available, the HMM could have been trained using the Baum-Welch algorithm (BAUM 1972) directly on individual MPP sample data. We have not tested this strategy but expect that it would require carefully chosen initial values of the emission distribution parameters and many MPP samples to ensure that we observed all possible genotypes at each gene locus.

We implemented the emission model as a multivariate normal distribution in which, conditional on the genotype state, the sample profile components are uncorrelated normal random variables. There are two challenges here. First, it required setting the scale (σ^$^) of the normal distributions. Second, it does not allow for differences in precision between profiles from highly versus lowly expressed genes. In principle we could have implemented a count-based model, e.g., a Dirichlet distribution, to circumvent both problems. However, the simpler model performs well, and we do not expect significant improvement would be achieved using a count model.

We implemented GBRS for DO mouse populations using the transmission model implemented in R/qtl2 (BROMAN et al., 2019). In our experience, HMMs for genotype reconstruction are robust to the magnitude of the transition probabilities. However, high transition rates can result in ‘choppy’ genome reconstructions with presumably false-positive recombination breakpoints, and low transition rates could result in ‘smoothing over’ of small haplotype segments. Results are best when the transition model is matched to the breeding design of the MPP. We have successfully applied GBRS to collaborative cross (CC) inbred strains, CC F1 hybrid mice, and a DO x C57BL/6 backcross by replacing the HMM transmission model. Implementing GBRS for other mouse models, e.g. UM-HET3 or BxD recombinant inbred strains, appears feasible but would also require founder expression data and retraining the emission model.

An advantage of working with mouse MPPs is that the genomic data resources (reference genome, genome annotation, and variant calls for founder strains) are readily available and highly accurate. We have described GBRS in general terms here because it could in principle be applied to any MPP, including non-rodent species. The documentation and open source GBRS software provide sufficient information to undertake implementation, but it would require a dedicated effort and remains untested.

Our implementation of GBRS, despite several shortcomings noted above, has demonstrated that there is sufficient genotype information in RNA-Seq reads from the DO mouse population to support practical applications. GBRS performs as well as array-based methods at both genome reconstruction and RNA-Seq quantification and it could be used as a stand-alone genotyping strategy. When used in conjunction with other genotyping methods, GBRS provides a highly effective quality control tool to detect and correct sample mix-ups and contamination and it can provide a means to recover samples with missing genotypes. GBRS as an adjunct to other genotyping methods provides a high level of confidence in genotyping accuracy and sample identity, as well as a simple, robust strategy to obtain total and allele-specific expression estimates that account for individual genetic variation.

## DATA AVAILABILITY

Founder data for the six tissues are accessioned in GEO: adipose GSE266920, gastrocnemius GSE266922, heart GSE266918, liver GSE266923, and pancreatic islet GSE266921. DO mouse data for five tissues are also accessioned in GEO adipose GSE266549, gastrocnemius GSE266567, heart GSE267727, liver GSE266569, and pancreatic islet GSE267728.

## SOFTWARE AVAILABILITY

We have implemented the GBRS algorithm in an open-source python package available at https://github.com/churchill-lab/gbrs with MIT license. The package is containerized and available at the Quay.io hub: https://quay.io/repository/jaxcompsci/gbrs_py3. Nextflow pipelines for the analysis of data with GBRS and the generation of inputs for non-DO populations are available at https://github.com/TheJacksonLaboratory/cs-nf-pipelines. For the DO, required data files are available at https://zenodo.org/records/8289936. R scripts for eQTL mapping are available at https://thejacksonlaboratory.github.io/Workflowr_Array_GBRS/index.html. A nextflow pipeline to run GBRS is available at https://github.com/TheJacksonLaboratory/cs-nf-pipelines. Instructions for executing GBRS from the command line are provided in the **Supplementary Material**.

## ACKNOWLEDGEMENT

We gratefully acknowledge the contribution of the Gene Expression Service at The Jackson Laboratory for expert assistance with the work described in this publication.

